# Predicting inbred parent synchrony at flowering for maize hybrid seed production by integrating crop growth model with whole genome prediction

**DOI:** 10.1101/2024.11.20.624538

**Authors:** Anabelle Laurent, Eugenia Munaro, Honghua Zhao, Frank Technow, Eric Whitted, Randy Clark, Juan Pablo San Martin, Radu Totir

## Abstract

One of the challenges of maize hybrid seed production is to ensure synchrony at flowering of the two inbred parents of a hybrid, which depends on the specific parental combination and environmental conditions of the production field. Maize flowering can be simulated using a mechanistic crop growth model that converts thermal time accumulation to leaf numbers based on inbred specific physiological parameter values. Heretofore, these inbred specific physiological parameters need to be measured or assigned based on prior knowledge. Here, we leverage genetic, environmental and management data to predict physiological parameters and simulate flowering phenotypes by using whole genome prediction methodology combined with a crop growth model (CGM-WGP) as part of in-field in-season inbred growth development. We use two estimation sets that differ in terms of management and weather information to test the robustness of our approach. As part of our findings, we demonstrate the importance of defining informative priors to generate biologically meaningful predictions of unobserved physiological parameters. Our CGM-WGP infrastructure is efficient at simulating flowering phenotypes. An important practical application of our method is the ability to recommend differential planting intervals for male and female maize inbreds used in commercial seed production fields to synchronize male and female flowering.

**Core ideas:**

- Synchrony at flowering of maize inbred parents is crucial for optimal pollination and consequently seed yield.
- Integrating WGP with CGM can accurately predict physiological parameters and simulate maize flowering phenotypes.
- CGM-WGP infrastructure can be used to optimize field operations for large scale maize hybrid seed production.

## 1. Introduction

In maize growth and development, flowering is a pivotal stage that marks the transition from the vegetative stages, characterized by the number of visible collared leaves, to the reproductive stage. The time elapsing between sowing and flowering is associated with the total number of leaves per plant (hereafter called *tln*) and the rate of leaf appearance (Tollenaar et al., 1979). Flowering time is typically scored as two dates, when 50% of the plants in a plot are shedding pollen and when 50% of the ears have visible extruded silks (Fan et al., 2022). Flowering can influence maize yield under abiotic stress conditions, such as drought (Bruce et al., 2002; Kim and Lee, 2023), which emphasizes the importance of optimized management of maize seed production fields.

Accurate predictions of in-field in-season maize inbred crop development are highly desirable for both seed product development and commercial seed production. They create the opportunity to improve management practices, be time-efficient, and optimize use of human labor (Worku et al., 2016; Michel et al., 2022; Fan et al., 2022). Specifically, one of the challenges of maize hybrid seed production is to ensure synchrony at flowering of the two inbred parents (*i.e.,* timing between male pollen shedding, also called anthesis, and female silking). Indeed, female organ stigmas are receptive for approximately five days after silking starts (Bassetti and Westgate, 1993; Anderson et al., 2004), and senescence happens eight days later if not pollinated. Synchrony at flowering can be addressed not only by planting delay between male and female inbred parents but also by clipping the two-to-four whorl leaves of the male at stages V4-V6 to delay pollen shed by two to three days, or by detasseling early to hasten silk emergence by one or two days (MacRobert et al., 2014). Furthermore, optimized synchrony at flowering has several additional operational benefits such as minimizing undesired outcrossing to ensure seed quality (Goggi et al., 2006) and decreasing production risks to ensure reliable commercial seed yield production. However, because of genetically determined differences in developmental rates of the inbred parents, their time to reach flowering can differ substantially. The development of the parents can also be affected differently by the environmental and management factors associated with the production fields they are growing in (Arisnabarreta and Solari, 2017). Thus, having a computational framework that makes use of genotype by environment by management interactions to accurately simulate female and male inbred development is desirable to optimize synchrony of flowering in practical field conditions.

Inbred maize flowering can be predicted using a mechanistic crop growth model (CGM) that converts thermal time accumulation (also called growing degree day, GDD) to leaf numbers by using physiological parameter values such as the leaf appearance rate (Wilhelm and McMaster, 1995) or phyllochron (Birch et al., 1998) like in the CGMs APSIM, DSSAT and CERES-Maize (Holzworth et al., 2014; Hoogenboom et al., 2019; Xu et al., 2023). In field conditions with no nutrient and water limitations, the relationship between the thermal time and the leaf appearance from emergence to flowering has been modelled with linear, exponential and bilinear mathematical functions (Muchow and Carberry, 1989; Birch et al., 1998; Abendroth et al., 2011; Dos Santos et al., 2022). Female silking and male shedding are associated to a series of genotype-specific physiological parameters, such as *tln*, that need to be measured or assigned to take full advantage of a CGM in hybrid seed production. This represents a major limitation for the utility of CGM as it is expensive and often not practical to measure genotype specific physiological parameters in field conditions (Boote et al., 1996; Messina et al., 2011; Bustos-Korts et al., 2019). Indeed, the measurement of flowering time requires regular visual evaluations of each plot during the flowering period, which is time-consuming and requires specialized domain knowledge (Fan et al., 2022). Another limitation of CGM is that simulations can be inaccurate if the physiological parameters are assigned to a constant value for all male and female inbreds as opposed to being genotype specific. For instance, genotype specific variability in the leaf appearance rate has been demonstrated by several authors (Muchow and Carberry, 1989; Padilla and Otegui, 2005; Van Esbroeck et al., 2008).

The whole genome prediction (WGP) set of statistical methods is a robust platform that can enable prediction of genotype specific physiological traits (Meuwissen et al., 2001) in active breeding programs. As commercial seed operations manage large portfolios of hybrids involving many inbred parents (Mikel and Dudley, 2006; White et al., 2020), obtaining accurate flowering time information for all the resulting parental combinations can be a challenge. Several scientific studies demonstrated that genomic prediction can be used for predicting maize flowering time (Li et al., 2018; Yuan et al., 2019; Wu et al., 2019; Fan et al., 2022). However, standard genomic prediction models are limited for prediction of flowering time for specific production locations, as required for optimizing inbred parent synchrony for hybrid seed production. In addition, WGP requires availability of data sets that can be challenging and expensive to construct for each physiological parameter needed by the CGM. To address this limitation, and thus take advantage of both CGM and WGP, Technow *et al*. (2015) proposed a novel methodology where CGMs are integrated with WGP through approximate Bayesian computation (CGM-WGP). Furthermore, to improve computational efficiency, the original approach of Technow et al., (2015) was extended to use Bayesian Hierarchical models (Cooper *et al*. 2016, 2017; Messina *et al*. 2018) and afterwards used to demonstrate its ability to generate accurate predictions of physiological parameters and accurate simulation of phenotypes (Onogi et al., 2016; Cooper et al., 2016, 2017; Messina et al., 2018; Campbell et al., 2020; Diepenbrock et al., 2022; Jighly et al., 2023).

Here we present an approach for predicting field specific maize inbred flowering time based on the above mentioned CGM-WGP methodology. The paper is organized as follows: i) we discuss technical aspects of the CGM-WGP, and the need to carefully specify informative priors to generate biologically meaningful predictions of unobserved physiological parameters, and ii) we discuss the use of CGM-WGP infrastructure for optimizing parent synchrony with an applied example from commercial seed production.

## 2. Materials & Methods

### 2.1 Description of the CGM-WGP infrastructure

We start by describing the crop growth model used in our study (see Figure 1) named Maize Flowering Synchrony model (hereafter called MFS). MFS is parametrized by three unobserved physiological parameters (*i.e*., coefficient of leaf appearance, hereafter called *coblf*, *tln,* and *ebR1*) that determine the phenotypes days-to-silk and days-to-shed as a function of thermal time accumulation (Figure 2). MFS is based on an exponential relationship between leaf number (LN) and thermal time (GDD) described by Muchow *et al*. (1990) where LN is equal to:

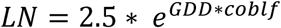

with 2.5 representing the number of leaves at emergence (Muchow and Carberry, 1989; Muchow et al., 1990), and *coblf* equal to 0.00225 in Muchow *et al*. (1990) but considered as a genotype-specific parameter in our study. After planting, 30.6 GDD (°C) are required for maize to germinate. Air temperature data was retrieved with the R package *nasapower* (Sparks, 2018; Sparks et al., 2023).

**Figure 1:**
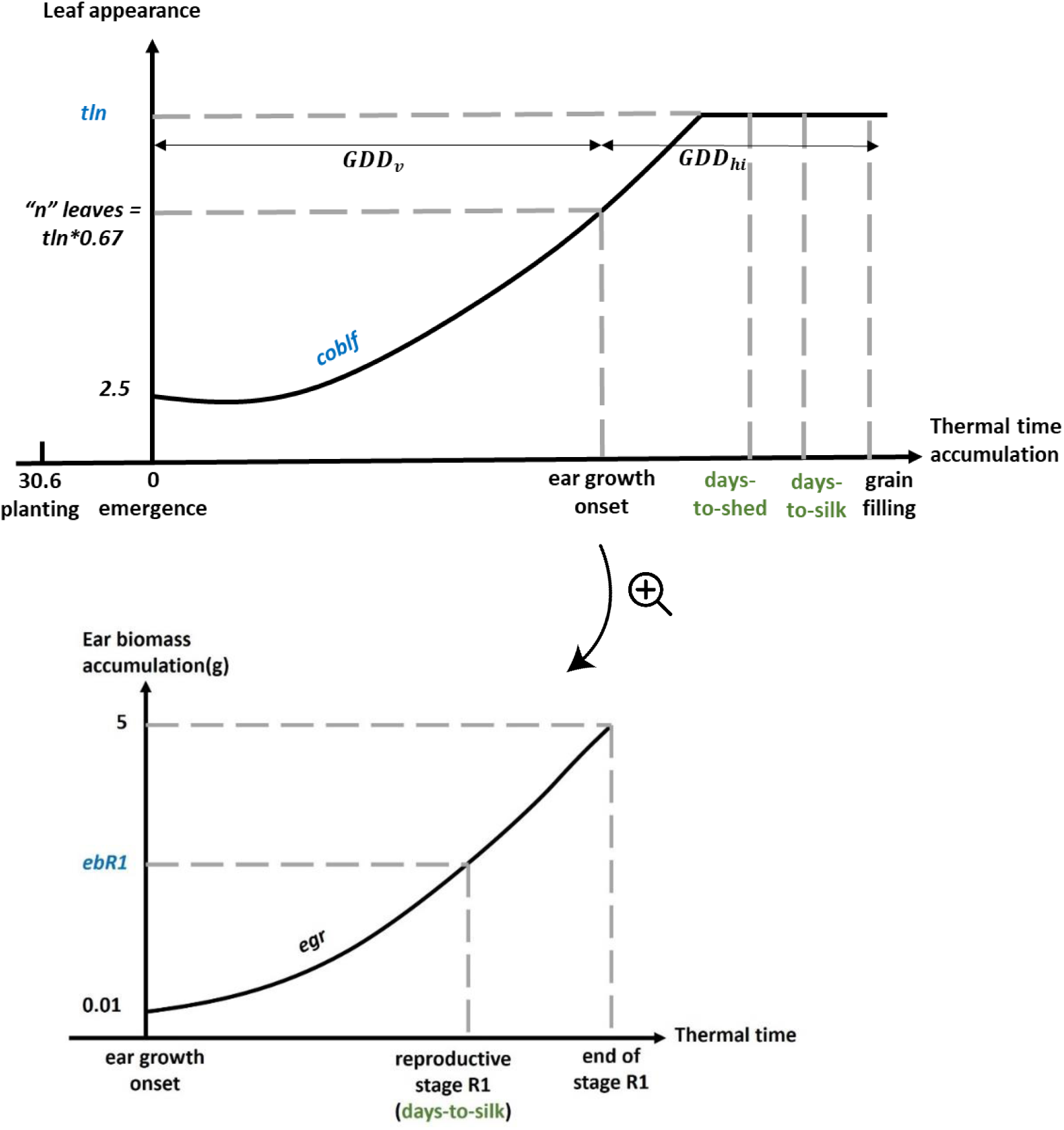
Description of the Maize Flowering Synchrony crop growth model. The three physiological parameters (*tln*, *coblf*, and *ebR1*) (in blue), determined the two phenotypes’ days-to-silk and days-to-shed (in green). The scheme at the top represents the relationship between leaf appearance and thermal time. The scheme at the bottom represents the ear biomass development over time.

**Figure 2:**
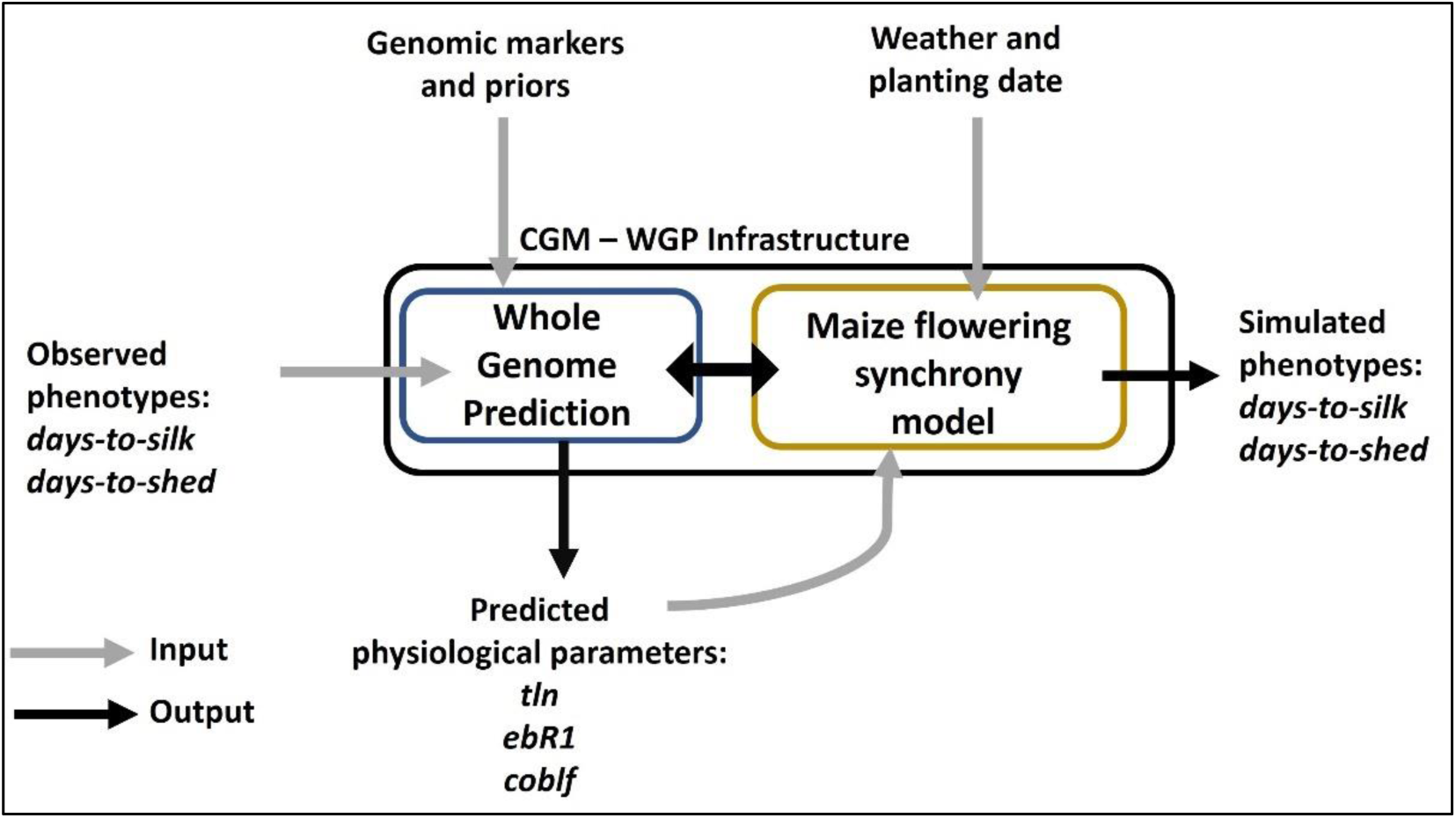
Description of the CGM-WGP infrastructure to simulate phenotypes for in-field in-season crop growth development. Our CGM is called Maize Flowering Synchrony.

After a specific thermal time accumulation *GDD_v_*, the onset of ear growth is initiated at a developmental stage “Vn”. It is assumed that the ear leaf coincides with the position of the largest leaf which is related to the parameter total leaf number (*tln*) as follow (Birch et al., 1998):

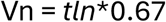

As an example, if a specific inbred has a total leaf number equal to 23, the ear growth will be initiated at the developmental stage “V15”.

To model female flowering (*i.e*., silking), an exponential growth pattern is used to calculate accumulated ear biomass (Cárcova et al., 2003; Borrás et al., 2007). The ear grows exponentially with an ear growth rate (*egr*) equal to 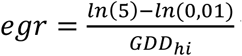 with *GDD* representing the thermal time between the development stage “Vn” to the beginning of grain filling as a function of *tln*. As a starting point of ear growth development, the ear biomass is considered equal to 0.01g and grows up to a maximum of 5g during the silking period (Otegui, 1997; Borrás et al., 2007; Cooper et al., 2014).

We consider 40 GDD (°C) between the expansion of the last leaf (i.e., *tln* is reached) and pollen shedding (refer as days-to-shed in Figure 1) (Muchow and Carberry, 1989).

Silking is a change of state at the individual plant level and when the ear biomass exceeds the value of *ebR1*, we consider the reproductive stage R1 (*i.e*., silking) reached. The ear biomass at a specific thermal time GDD is equal to 0.01 * *e^egr*GDD^*

The MFS crop growth model was incorporated into the CGM-WGP sampling algorithm presented by Messina *et al*. (2018) as described in Figure 2. It is important to note that the three physiological parameters *tln*, *coblf*, and *ebR1* are genotype specific and are modeled in CGM-WGP as a linear function of observed biallelic single nucleotide polymorphism (SNP) markers representing the inbred genotypes. Those physiological parameters are then connected to the observed flowering phenotypes by the MFS which serves as a link function.

In the context of the CGM-WGP infrastructure, the MFS crop growth model can be used as a link function between the physiological parameters and the observed phenotypic data as represented by the general equation below (1):

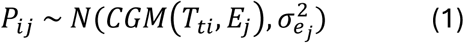

Where *P_ij_* is the simulated phenotype (in our study days-to-silk and days-to-shed) of the genotype *i* in the field *j*. *T_ti_* is the genotype-specific unobserved physiological trait *t* for the genotype *i*; *E_j_* represents the environmental features for the field *j* and *σ*^2^ is the residual variance for the simulated phenotype in the field *j.* The prior on the genotype-specific physiological trait *T_ti_* (*i.e*., coefficient of leaf appearance *coblf*, total leaf number *tln* and ear biomass at silking stage R1 *ebR1*) is:

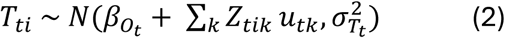

Where 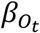 is the intercept of the genotype-specific trait *t*; *Z_tik_* is the SNP score of the genotype *i* at marker *k* of trait *t* (same markers were used for all traits); *u_tk_* is the effect of the marker *k* for trait *t* (same markers were used for all traits); 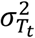 is prior variance for the trait *t*. The prior distribution for 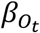 is a Normal distribution with mean 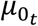 and a residual variance 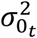. Priors for *μ_tk_* are Normal distributions with mean of 0 and marker effect variance of 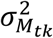, which is equivalent to those used in ‘BayesA’ WGP model (Meuwissen et al., 2001). The variances 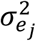, 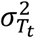 and 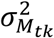 are associated with a scaled inverse chi-square prior distribution with 4.001 degrees of freedom and scaling factors *S_t_*. *S_t_* follows a gamma distributions with shape and rate parameters calculated according to Habier et al. (2011). This hierarchical model can be visualized in a tree form and more information are provided in Messina *et al*. (2018).

Metropolis-Hastings within Gibbs algorithm was implemented to sample from the posterior distribution of all the parameters (Gelman, 2004). Sampling chains were run for 50,000 iterations with the first 30,000 discarded as ‘burn-in’ and samples from every 20^th^ postburn-in iteration stored, thus resulting in 1000 stored samples. The algorithm was implemented as a C++ routine.

The posterior predictive distribution for a genotype *i* in field j was obtained by entering the physiological parameters *T_ti_* into the MFS model together with the inputs of the field *j*, resulting in one set of phenotypic values per posterior sample. The mean of this distribution is used as the predicted value. Note that the predicted values of the set of physiological parameters are used as a joint set in the MFS model to describe the relationship between thermal time and inbred development. Indeed, following the WGP step, the set of new physiological parameters sampled from the posterior distribution, if accepted, are passed as a joint set to the MFS model together with the management and weather information (See Figure 2) to simulate the genotype specific flowering phenotypes. However, jointly sampling the set of physiological parameters as described above can create a non-identifiability problem, meaning that different combinations for the three physiological parameters can lead to the same phenotype value (Wechsler et al., 2013).

### 2.2 Data collection

We collected data from three maize fields in the US Corn Belt (Iowa, IA) from the years 2018, 2020, and 2021. As each field is composed of multiple plots, a given inbred can be grown multiple times and thus have several observed phenotypes. When 50% of the plants in each plot were shedding pollen (for males) and silking (for females), the corresponding dates were recorded and converted in number of days from the planting date to the flowering date.

Regarding the summaries of weather data, 2018 had high GDD while 2020 had the lowest GDD accumulation (Figure 3). We provided additional weather characteristics in the Supplemental Material (section S1).

**Figure 3:**
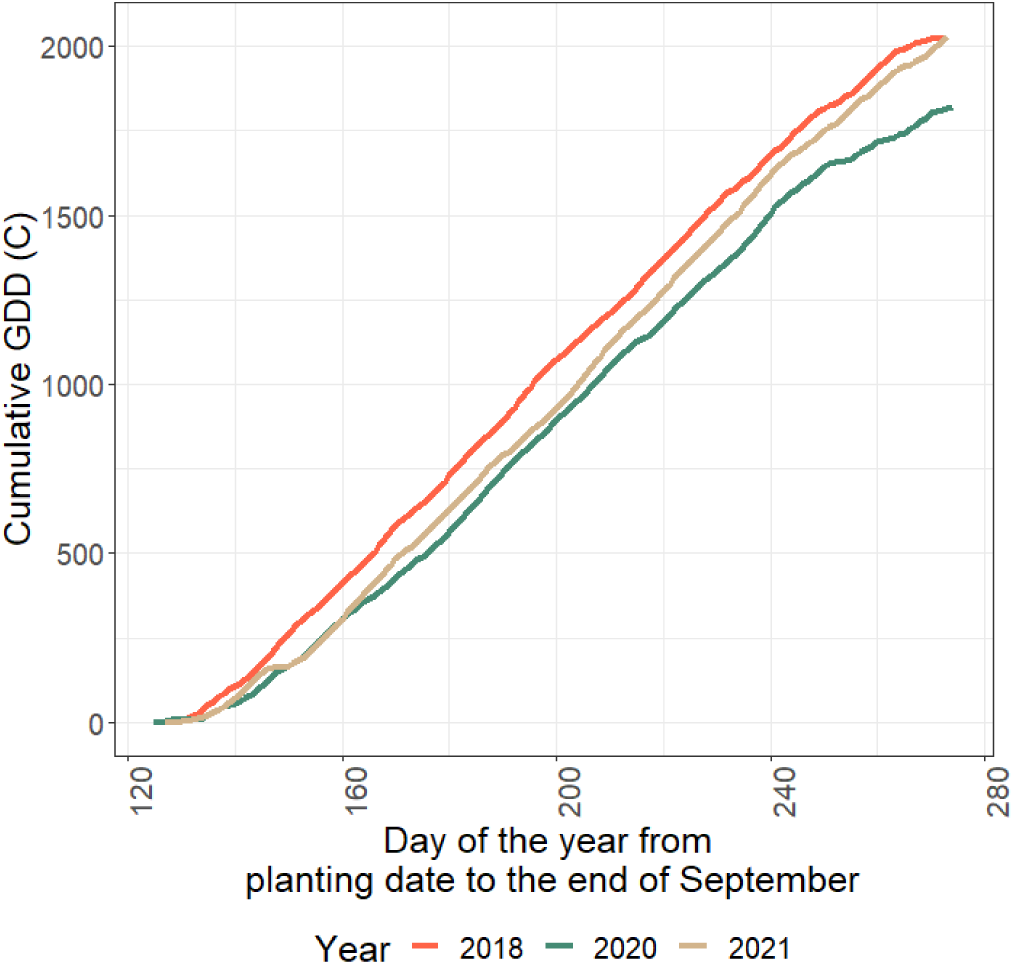
Cumulative GDD-Celsius from planting date to September 30^th^ for the three fields conducted in 2018 (red), 2020 (green), and 2021 (beige).

We also used a physiological parameter validation set, where the total leaf numbers were measured for 26 genotypes grown in 2021 in fields from both Iowa and Nebraska (Figure 4). The protocol to count the total leaf number was as follow: when reaching the developmental stage V4 at the field scale, the number of leaves was counted for four representative plants per plot (*i.e.*, per genotype) using the leaf collar method (Licht, 2023). The representative plants were tagged for future observations and by marking the topmost collared leaf. The observations are repeated every 10 days and only the new leaves after the marked leaf were counted.

**Figure 4:**
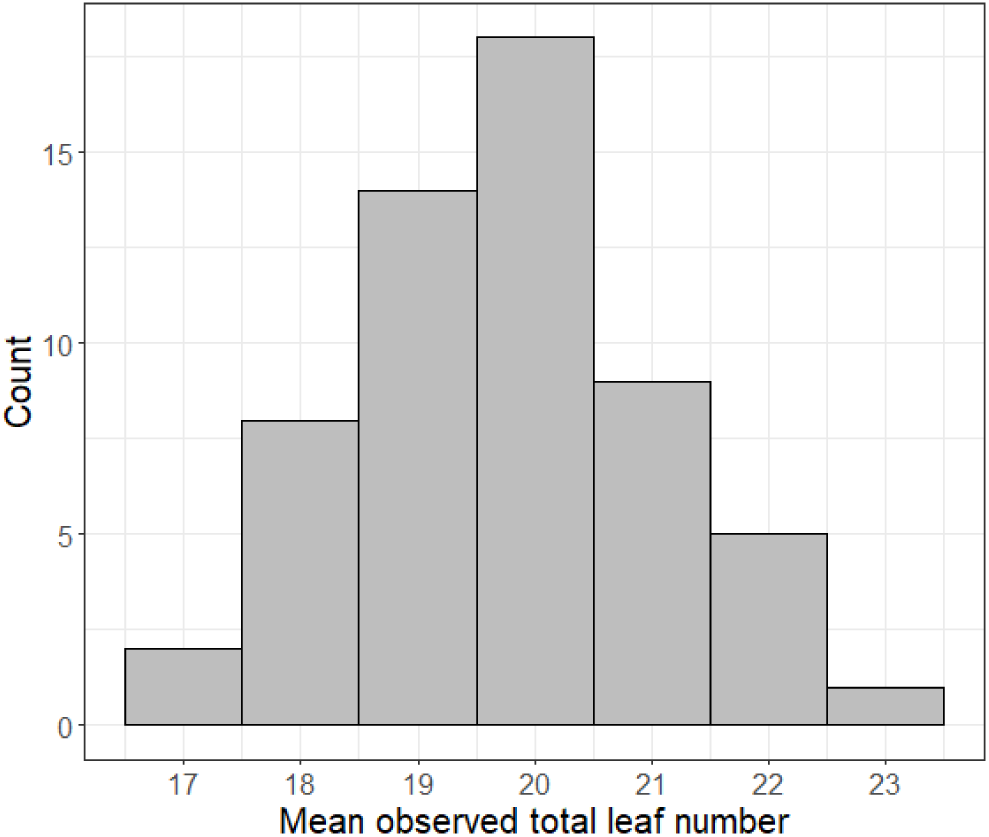
Distribution of the mean observed total leaf number across 26 genotypes evaluated in 2021 in Nebraska and Iowa. The mean is calculated for each genotype with a maximum of four samples.

### 2.3 Data sets description

We used a data set (called data set A) with data collected from three maize fields described above. Data set A comprises of ∼2000 female inbreds. We also used a second data set (called data set B), built as a subset of data set A, which includes ∼500 female inbreds collected from one field conducted in 2021.

We used a data set (called data set C) which comprises of ∼2000 male inbreds with data collected from three maize fields described above.

The MFS-CGM was fitted separately for the data sets A, B, and C.

### 2.4 Definition of Bayesian priors for the physiological parameters

As our intent is to assess the impact of the Bayesian priors on the quality of the predictions for the physiological parameters *T_ti_* and the simulations of the phenotypes, we explored two different priors for *ebR1* only while the priors for *tln* and *coblf* remained unchanged. As a reminder, the MFS model considers that the reproductive stage R1 is reached when the ear biomass accumulation exceeds a specific weight representing the ear biomass at which 50% of the plants have visible silks (Figure 1). To be in accordance with our crop growth model, we defined a prior (called *ebR1_50*) that follows a Normal distribution centered on 2 and with a standard deviation of 0.5 (Figures 5 and 6.B) (Khan et al., 2022). Another prior mean for *ebR1* was defined to represent the ear biomass at the beginning of the silk appearance (i.e., when the ear has its very first visible silk, which can be associated at 1% of the silk appearance period, see Figure 5 in orange and Figure 6A). This prior mean, called *ebR1_1*, is based on a study from Borrás and Vitantonio-Mazzini (2018) and follows a Normal distribution centered on 0.56 and with a standard deviation of 0.089 (Figure 6.A).

**Figure 5:**
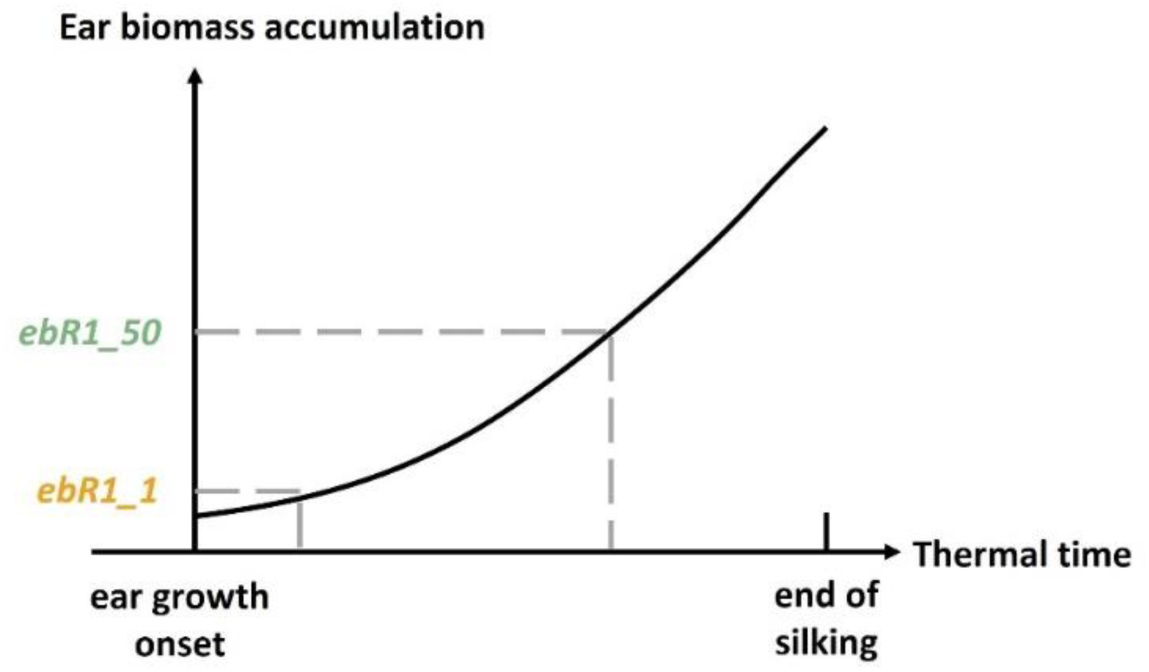
Description of the two Bayesian priors tested to represent the intercept of the physiological parameter *ebR1*.

**Figure 6:**
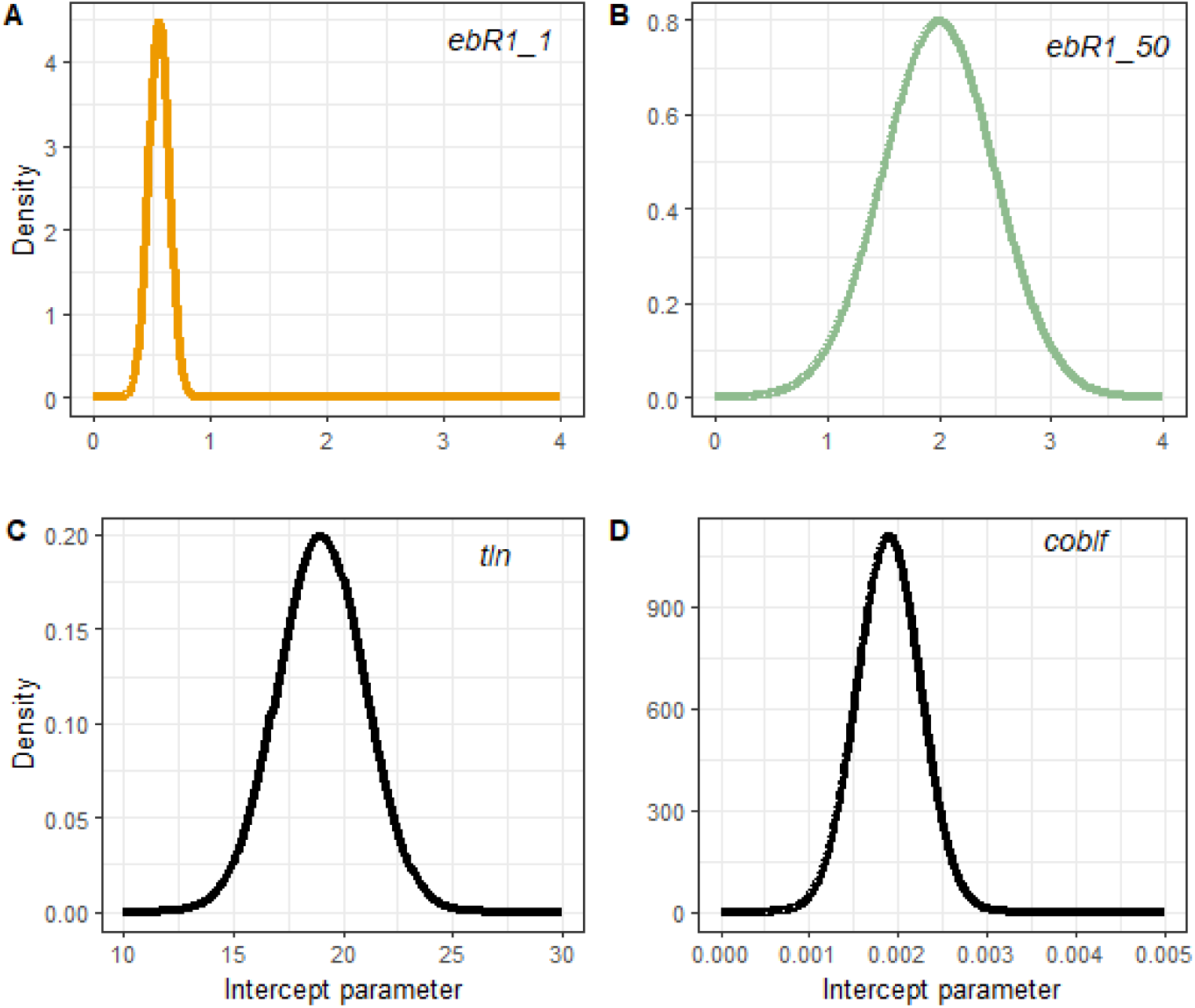
Normal distribution used for the intercept of the parameters ebR1 (A and B, ebR1_1 and ebR1_50, respectively), *tln* (C) and *coblf* (D). *ebR1_1*∼ N(0.56, 0.0089); *ebR1_50*∼ N(2, 0.5); *tln*∼ N(19, 2); *coblf*∼ N(0.0019, 0.00036).

The physiological parameter *tln* follows a Normal distribution centered on 19 with a standard deviation of 2, *coblf* follows a Normal distribution centered on 0.0019 with a standard deviation of 0.00036 (Muchow and Carberry, 1989; Muchow et al., 1990) (Figure 6 C-D).

We also defined uninformative priors that follow a Normal distribution with the same mean for for *ebR1_50*, *tln* and *coblf* but with a larger standard deviation. This can be seen as baseline priors. A more detailed description of the priors and results of the impact of uninformative priors to the simulated phenotypes are shared in Supplemental Materials (section S2).

### 2.4 Measure of accuracy and prediction quality

The Pearson correlation coefficient, the mean absolute error (MAE) between the simulated and the measured phenotype (days-to-silk), and the root mean square error (RMSE) were calculated to assess the goodness of fit of MFS-WGP.

### 2.5 Optimizing seed operations by simulating maize flowering synchrony

We demonstrate the value of using genotype specific inbred parent phenotypes for optimizing large scale field operations for seed production, with a real-life application example. From the data set A, we selected one female inbred, called Fem1, and from the data set C, we selected two male inbreds, called Mal1 and Mal2. They were all grown in three fields and each of their phenotypes was measured. The MFS-WGP was fitted separately for the data set A but excluding the inbred parent Fem1 and for the data set C excluding the inbred patents Mal1 and Mal2.

Two maize hybrids, called Hyb1 and Hyb2, were created by crossing the inbred parents Fem1xMal1 and Fem1xMal2, respectively.

## 3. Results

### 3.1 Predictive ability

MFS-WGP is able to predict flowering in a different year, and the Pearson correlation coefficient shows no major difference regardless of the size of the estimation set (Table 1).

**Table 1:** Predictive ability of days-to-silk of female inbreds (defined here as Pearson correlation coefficient between the vector of MFS-WGP simulations and the vector of measured phenotypes).

| **Estimation set** | **Validation set** | **CGM-WGP, Pearson correlation coefficient** |
| --- | --- | --- |
| 2018 | 2021 | 0.78 |
| 2020 | 2021 | 0.80 |
| 2018-2020 | 2021 | 0.81 |

### 3.2 Importance of specification of Bayesian priors to obtain biologically meaningful predictions for physiological parameters

When the data set is large and diverse in terms of fields and genotypes (data set A), the relationship between the simulated and observed days-to-silk is similar whatever the prior on the intercept of *ebR1*. Indeed, the MAE is equal to 0.68 and 0.66, and RMSE equal to 0.88 and 0.85, for *ebR1_1* and *ebR1_50*, respectively (Figure 7). On the other hand, when the data set is small and restricted to one field and only ∼16% of the genotypes (data set B), there is a strong impact of the prior chosen as shown by the MAE and the RMSE (MAE equals 0.02 and 0.59, and RMSE equals to 0.07 and 0.81, for *ebR1_1* and *ebR1_50*, respectively). In addition, as data set B includes only one field and one year the range of values for the observed days-to-silk is limited (from 65 to 79) compared to the data set A (from 54 to 84) (Figure 7).

**Figure 7:**
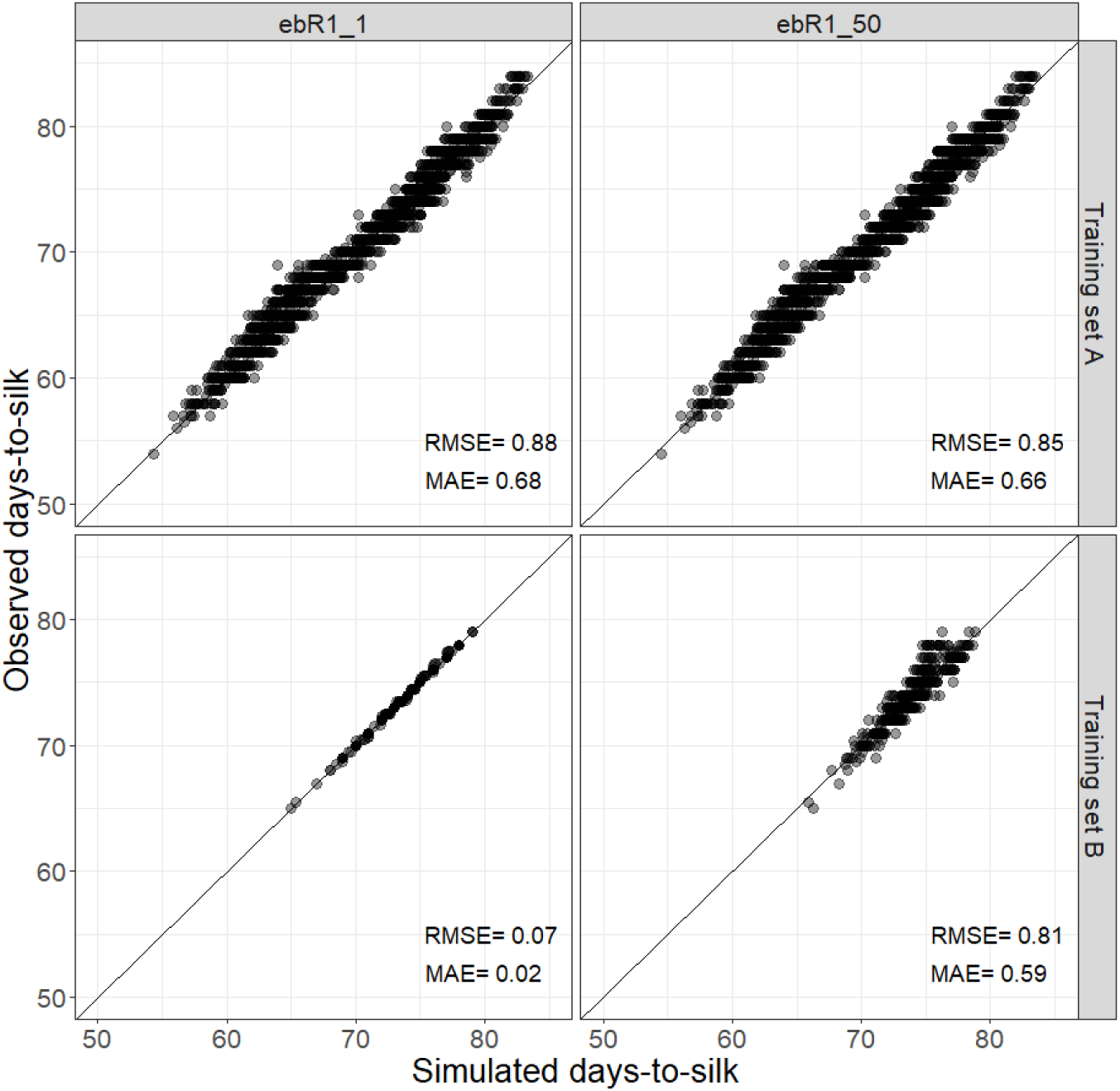
Simulated days-to-silk compared to the observed value by data set A (large data set of ∼2000 genotypes, at the top) and data set B (smaller data set of ∼500 genotypes, at the bottom) for the two types of priors for the intercept of *ebR1* (*ebR1_1* on the left, *ebR1_50 on the right*). The mean absolute error (MAE) and root mean square error (RMSE) are indicated at the bottom right of each plot. The identity line is indicated in black.

The posterior distributions of the intercept of 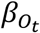 of *tln* and *ebR1* and the distribution of the posterior mean of *tln* and *ebR1* (Figures 8-9) help understand the simulated days-to-silk returned by the MFS-WGP infrastructure. For the small data set B, it is observed that the posterior distribution of the intercept 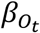 for *tln* has small variability for ebR1_1 (22±0.5 leaves) but not for *ebR1_50* (19-22 leaves) (Figure 8A). For the large data set A, the posterior distribution of the intercept 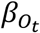 for *tln* has small variability regardless the value of *ebR1* (Figure 8C). The posterior distribution of the prior mean for total leaf number for the data set B with *ebR1_50* prior has most of its density between 18 and 22 leaves while with *ebR1_1*, the density covers a large range of value from 18 to 25 (Figure 8D). For the large data set A, the posterior distribution of the prior mean for total leaf number is similar between *ebR1_1* and *ebR1_50* (Figure 8D).The grey boxplot that represents the distribution of observed *tln* for 26 genotypes falls better within the posterior distribution of the prior mean of *tln* with the prior ebR1_50 (Figure 8 C-D).

**Figure 8:**
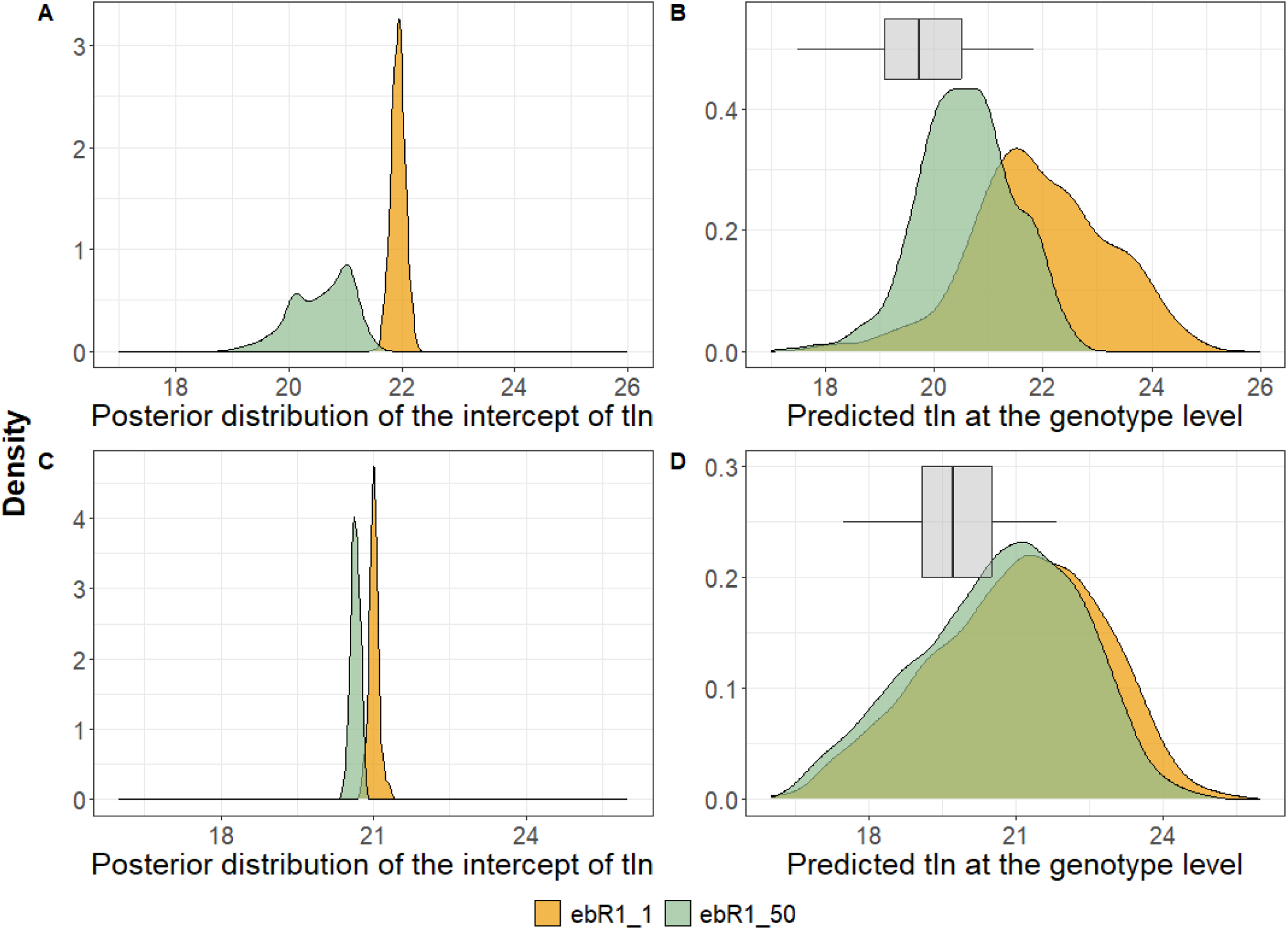
Impact of the priors *ebR1_1* (in yellow) and *ebR1_50* (in green) on the posterior distribution of the intercept 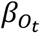 and on the distribution of the posterior mean of *T_ti_* for *tln* of the ∼500 female lines in the data set B (A) (B) and of the ∼2000 female lines in the data set A (C) (D). The grey boxplot (C) represents the distribution of observed total leaf number for 26 genotypes. The x-axis is in number of leaves.

**Figure 9:**
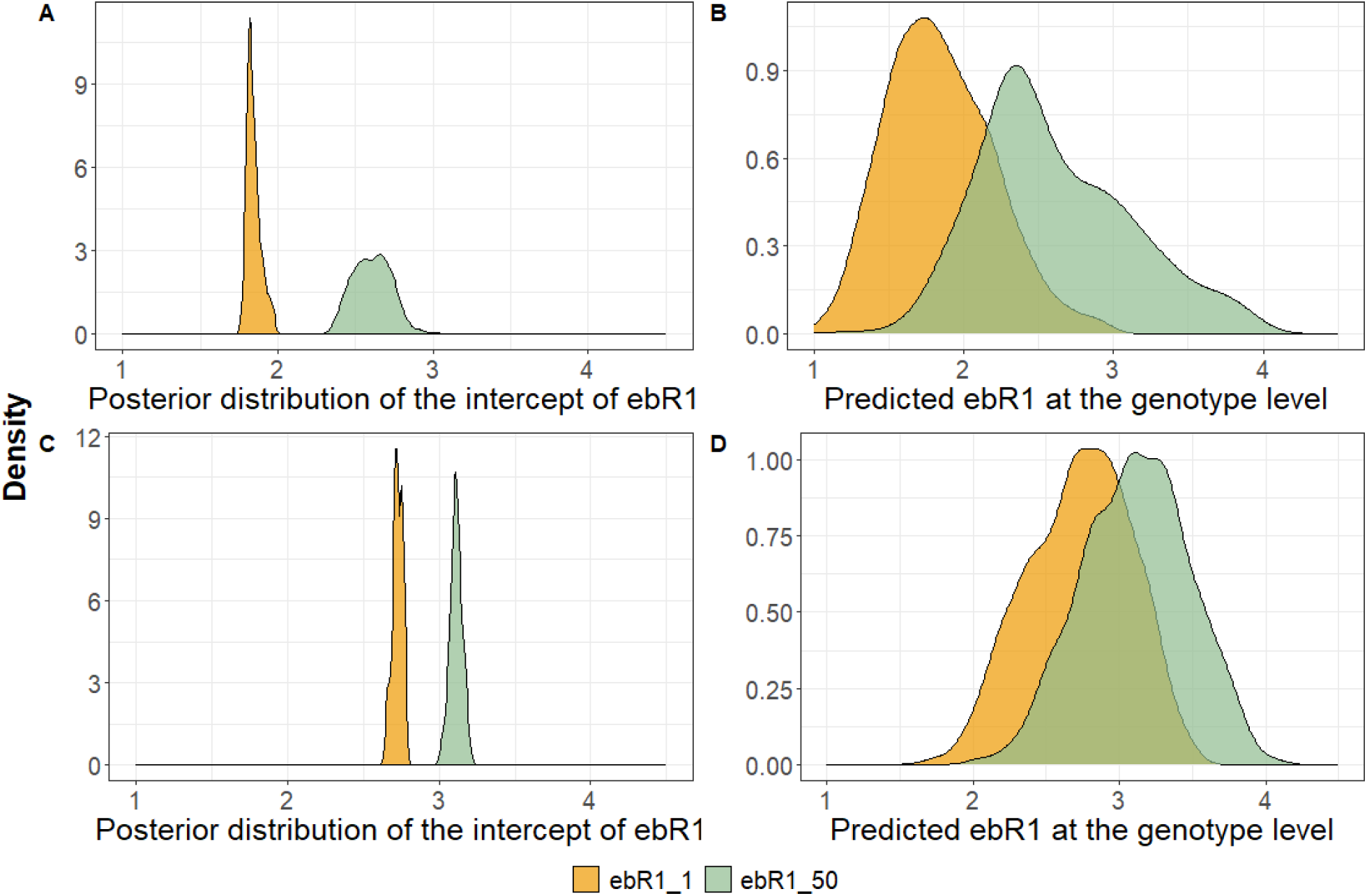
Impact of the priors *ebR1_1* (in yellow) and *ebR1_50* (in green) on the posterior distribution of the intercept 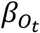 and on the distribution of the posterior mean of *T_ti_* for *ebR1* of the ∼500 female lines in the data set B (A) (B) and of the ∼2000 female lines in the data set A (C) (D). The x-axis is in grams.

For the small data set B, the posterior distribution of the intercept 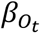for *ebR1* is centered around 1.8g for ebR1_1 and around 2.6g for ebR1_50 (Figure 9A) while for the data set A, the distribution are closer to 3g (Figure 9 C). For the data set B, the posterior distribution of the prior mean for ear biomass at silking is shifted to the left (ranging from 1 to 3) for the *ebR1_1* compared to *ebR1_50* (ranging from 1.5 to 4) (Figure 9B). For the data set A, the posterior distribution of the prior mean for ear biomass at silking for *ebR1_1* and *ebR1_50* overlapped more between them but with a slight shift to the left for *ebR1_1* (Figure 9D).

A 3D visualization describes the relationship between the three physiological parameters and the days-to-silk value (Figure 10) and illustrates that several combinations of the three physiological parameters result in similar days-to-silk.

**Figure 10:**
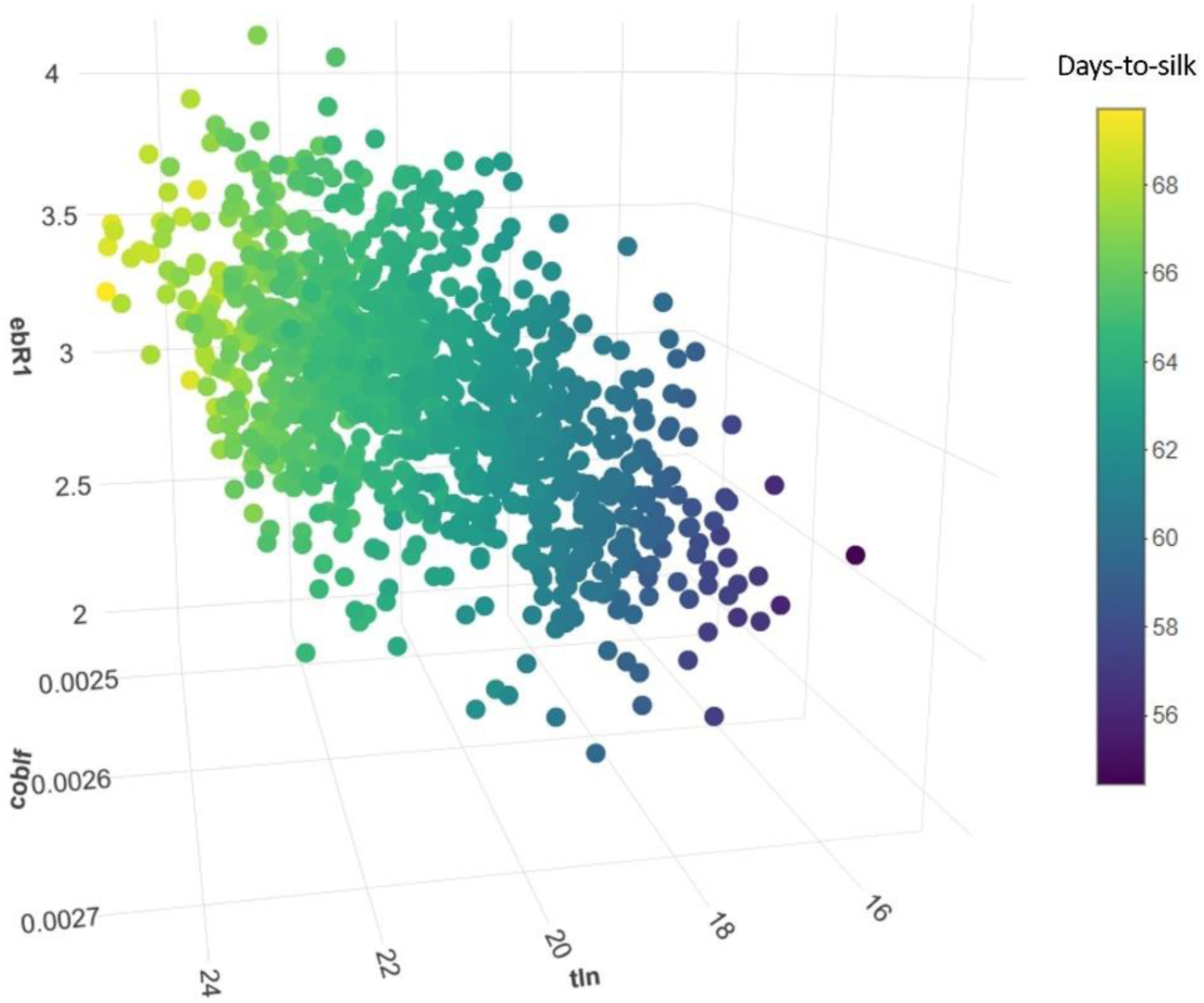
3D visualization representing the relationship between the three physiological parameters (x, y and z axis) and the phenotype day-to-silk (indicated in color) for the data set A.

### 3.3 In-field in-season crop growth development and flowering synchrony

The difference between days-to-silk for the female Fem1 and days-to-shed for the two males (Mal1 and Mal2) (also called anthesis silking interval), when measured or simulated with MFS-WGP, is displayed in Figure 11 for three different fields. When producing commercial seed for the hybrid Hyb1, because its two parents have similar relative maturities and flowering dynamics, both the observed and simulated difference between days-to-silk for female Fem1 and days-to-shed for male Mal1 are synchronous (Figure 11 at the top). Because of the synchrony in flowering between the two parents, no differential planting is needed for the two parents to ensure optimized hybrid seed production. However, when producing hybrid Hyb2, chosen here as a contrasting example, the two parents have an average difference between days-to-silk for female Fem1 and days-to-shed for male Mal2 of approximately five days (Figure 11 lower panel).

**Figure 11:**
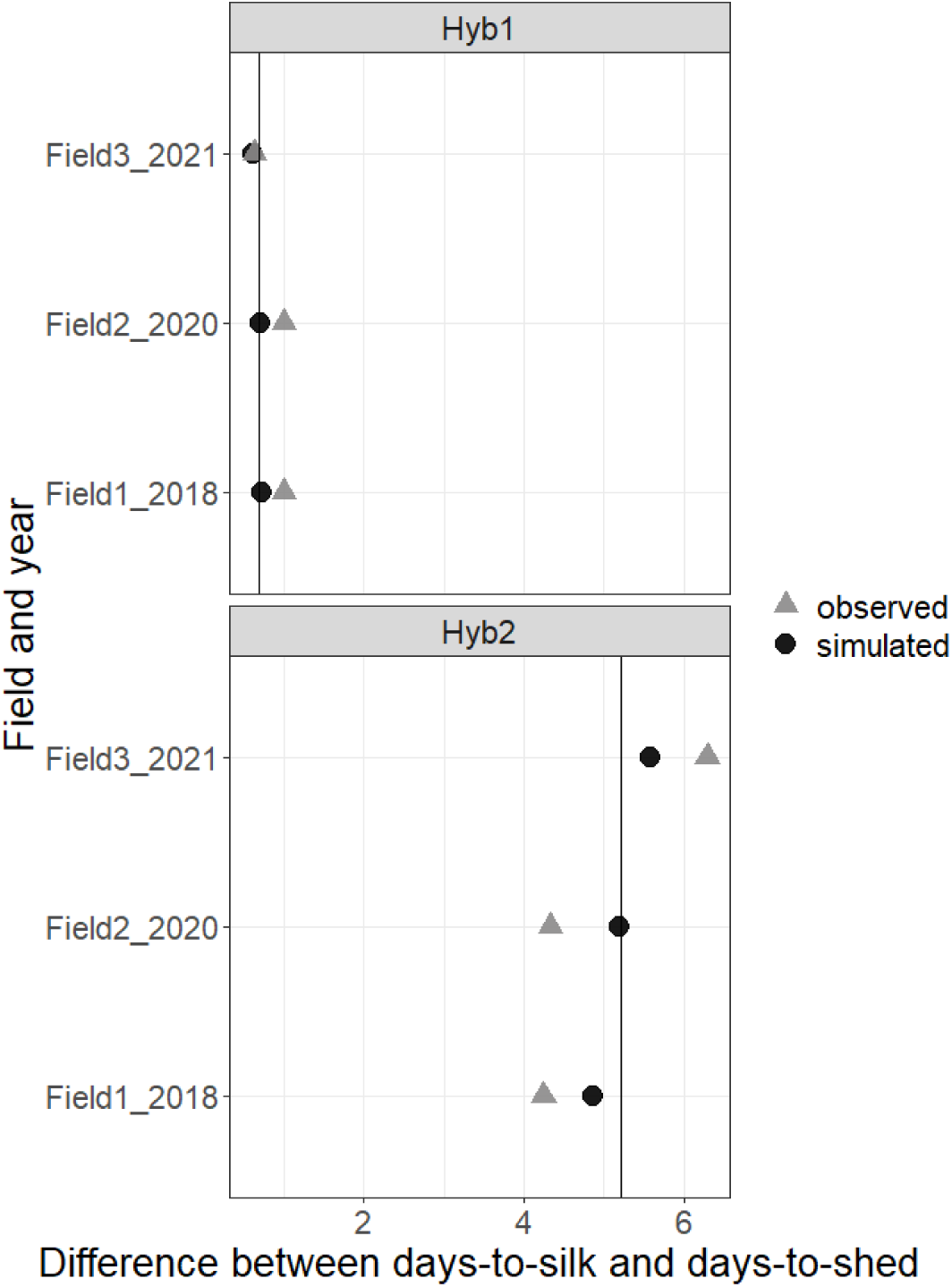
Difference between the days-to-silk and days-to-shed (simulated and observed values are represented by a black dot and a grey triangle, respectively) for two different hybrids (Hyb1, Hyb2) in three fields conducted at different years. The vertical black line represents the mean of the simulated difference across the three fields.

To understand how differential in planting dates could vary in hybrid seed production, a more general representation of the difference between days-to-silk and days-to-shed for a possible complete set of 2500 hybrids is represented in Figure 12. Those hybrids represent all possible crosses between 50 females and 50 males selected from data set A. Note that all hybrids represented in Figure 12 would not exist in hybrid seed production due to poor genetic and/or yield performance, but our point here is to illustrate the range in the difference between days-to-silk and days-to-shed. Indeed, the difference between days-to-silk and days-to-shed varies from −11 to 11 days. For any given male inbred (any y-axis tick on Figure 12), its planting date in commercial seed production fields would be impacted by the choice of the female inbred. The difference between days-to-silk and days-to-shed is comparable over the three locations.

**Figure 12:**
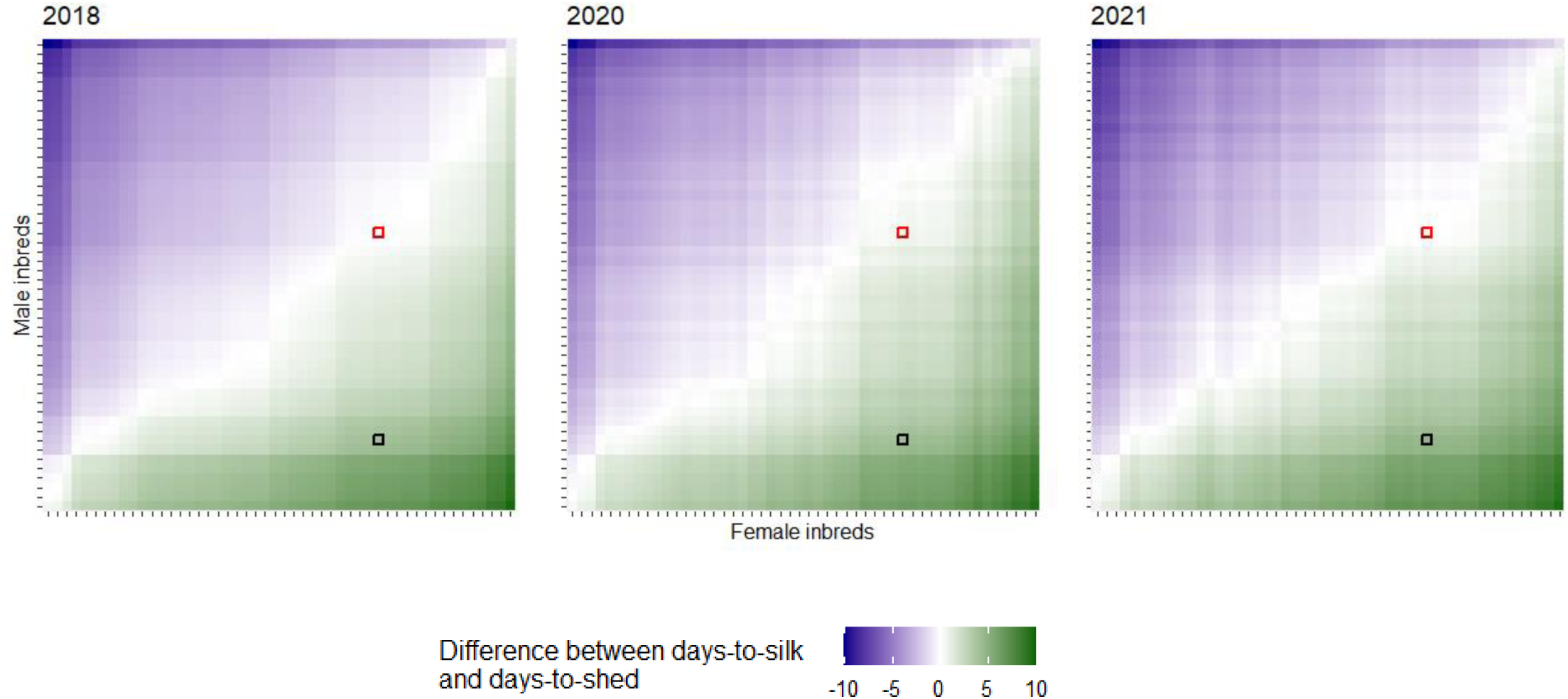
Difference between genotype specific simulated days-to-silk and simulated days-to-shed for a set of 50 female and male inbreds (x-axis and y-axis) based on 2018, 2020 and 2021 weather data. Females and males inbreds are ranked by increasing value of days-to-silk and days-to-shed using the location 2018 and the order is kept the same for 2020 and 2021. The hybrids Hyb1 and Hyb2, from figure 11, are represented by a red and black tile, respectively.

## 4. Discussion

The present study demonstrates the effectiveness of WGP combined with the MFS crop growth model to predict genotype-specific physiological parameters and simulate flowering phenotypes as part of in-field in-season inbred growth development. We show the importance of specifying biologically meaningful priors to generate interpretable predictions of the physiological parameters. In this context, our approach shows robustness, regardless of the characteristics of the estimation set (number of fields, growing seasons, and inbreds)

It is a common practice to calibrate cultivar parameters to simulate crop development in CGM (Jones et al., 2011). <u>Hoogenboom et al. (1999)</u> outlined a step-wise method for calibrating the developmental timing of a cultivar and the corresponding parameters for each step in an attempt to establish expert system rules to follow. Because the MFS can generate the same phenotype for different sets of physiological parameters, non-identifiability can be a challenge. As we have seen, the simulated phenotype can still be biologically plausible while the values of the physiological parameters may not be. Similar in objective to step-wise calibration method, the use of informative priors that are biologically plausible and meaningful within CGM-WGP framework also aims at overcoming lack of parameter identifiability that produces the expected simulated phenotype despite wrong physiological parameter values. In our study, in case of a prior misspecification (*ebR1_1*), especially with a small data set, the relationship between *tln* and ear ontogeny, as defined in MFS, results in compensations among processes that render estimated values of *tln* that do not align with observed values. The thermal time needed to reach flowering is achieved by increasing *tln* values beyond those observed because of the restriction on ear biomass growth imposed by the prior misspecification of *ebR1_1,* and consequently delaying the start of ear growth. As a result, the timing of vegetative stages and start of ear growth would be incorrect, limiting the use of the MFS as a decision tool that could aid in making critical decisions related to seed production such as irrigation, fertilizer and herbicide applications.

We have successfully demonstrated that non-identifiability at the physiological trait level can be overcome by using carefully chosen informative priors that are biologically meaningful. In addition, the data set should be as diverse and large as possible in terms of environments and individual inbreds to constrain the possible solutions for the genotype-specific physiological parameters and thus eliminate the non-identifiability issue (Poudel et al., 2023). In our case, increasing the diversity and informativeness of our data set (comparing data set A and the smaller and more narrow data set B) reduced the non-identifiability problem and improved the biological relevance of the prediction of the physiological parameters even in a case of prior misspecification (*ebR1_1*).

Unlike previous studies focused on predicting potential kernel production (Fonseca et al., 2004) or outcrossing in hybrid maize production (Astini et al., 2009) for which modeling flowering dynamics (*i.e*., amount and temporal distribution of exerted silks and pollen shed) of inbred parents is a key feature, the MFS is concerned with capturing the impact of differential planting dates between female and male inbreds on ASI, with no regards to specific parameters related to pollen production, pattern of silk emergence and pollination. Of all the genetic and management variables tested in these studies, however, altering ASIs had the greatest impact on female yield (Fonseca et al.,2004; Chassaigne-Ricciulli *et al*.,2021). Moreover, models of flowering dynamics require considerably more parameters and translating qualitative descriptive terms used in seed industry to characterize an inbred (e.g. “good”, “fair” and “poor” for male inbred pollen shed) into quantitative parameters can be challenging. The scope of applicability of this type of models is restricted by the correct parametrization and data availability, a limitation recognized by the authors.

The CGM-WGP framework described here strikes a balance between simplicity and accuracy of prediction of male and female flowering enabling its broad application as a decision tool in seed field management. Because of its simplicity in capturing main determinants of ASI in non-stress conditions and coupled with WGP in estimating genotype specific parameters it lends itself to be used for a large set of inbreds included in a breeding program. One of the main practical aspects of this work is to simulate the impact of differential planting dates between female and male inbreds used in commercial seed production. The ability to do so is crucial to ensure synchrony at flowering for optimal pollination which directly impacts seed set and consequently seed yield. Having the designated male release pollen at the right time also decreases the risk of hybrid seed lot contamination with external pollen, thus ensuring quality and purity of seed produced at commercial scale. Viable pollen is released for only six days after shedding starts (Uribelarrea et al., 2002; Westgate et al., 2003; Chassaigne-Ricciulli et al., 2021) which emphasize the need of being release at the right time. When the difference between the simulated days-to-silk and days-to shed is less or equal to two days, a differential of planting date between male and female inbred parents is not necessary. When the difference is higher than two days between the simulated days-to-silk and days-to-shed, differential planting dates must be considered to ensure reliable hybrid seed production. From the example of hybrid Hyb2 (Figure 11), since the male inbred is on average five days earlier than the female inbred, one could think that to have synchrony at flowering, the male inbred should be planted five days after the planting date of the female inbred. However, to optimize the chance of effective release of pollen at the period of exposure of the stigmas of the female, the male should be shedding two days prior and two days after the female starts silking. Thus, in this example, the male should be planted at two different dates: three and seven days after the female planting date. For both hybrids, the differences between the simulated, when removed from the data set, male and female flowering phenotypes are in accordance with the observed values. There is a shrinkage effect towards the simulated mean difference between the two phenotypes. As a reminder, those simulated days-to-silk and days-to-shed are direct outputs from the MFS crop growth model that simulates the growth and development as a function of thermal time accumulated (Figure 1). Therefore, in addition to the flowering dates, the vegetative stages can be simulated which is beneficial for the management of the field. Our mathematical modelling-based approach is not limited only to simulating inbred parent maize synchrony at flowering but can also be used to simulate other genotype-specific maize inbred development patterns using accurate forward looking weather forecasts. This could be highly beneficial for timing the application of fertilizer or pesticides or anticipating harvest dates, thus helping to optimize the logistics of commercial hybrid seed production. In terms of machinery and technical crews, deployment to perform field activities such as sampling, monitoring, and scouting.

The CGM-WGP approach involves estimation of unobserved, latent physiological parameters at various hierarchical levels, connected with each other and the output phenotypes as well as the environmental inputs through non-linear equations formulated in the CGM. This is, in principle, similar to the structure of machine learning methods such as neural networks, which are under active exploration and development for applications in breeding and agronomy (Fernandes et al., 2024; Crossa et al., 2024). The key difference, however, is that in CGM-WGP, these latent physiological parameters and connecting equations are informed and constrained by biological knowledge accumulated through often decades worth of research in crop biology and physiology (Hammer et al., 2006). One practical advantage is considerably less data is required for fitting robust CGM-WGP models, because the latent parameters and non-linear equations are specified as a priori and do not need to be learned from data. This also allows CGM-WGP to extrapolate beyond the observation space, if the underlying biological processes don’t change substantially (Hammer et al., 2006). Furthermore, the estimated latent parameters are readily biologically interpretable, which not only acts as a safeguard against overfitting and estimation errors (*i.e*., when parameters are estimated to be outside the biologically plausible range), but also allows to formulate novel biological hypothesis and targets for indirect selection. Nonetheless, agnostic, unsupervised non-linear machine learning methods are an alternative worth exploring, particularly for scenarios where sufficient prior biological knowledge is not available or where biological interpretation of parameters is not of interest.

The ability of our integrated CGM-WGP mathematical infrastructure to simulate genotype-specific in-field in-season inbred growth and development has significant advantages compared to the use of WGP or a CGM alone. At the scale of a breeding program, the WGP-CGM infrastructure can be used for predicting, evaluating, and selecting both resource-intensive and not readily scalable phenotypic component traits, such as total leaf number, coefficient of leaf appearance and ear biomass, as well as flowering traits for inbreds without phenotypic records measured in all environments. The prediction accuracy of the physiological parameters is nearly impossible to compute in a breeding program, except for very small data sets like the one used in our manuscript and for parameters such as *tln*. The rapid development of high-throughput phenotyping techniques such as image processing for leaf counting (Miao et al., 2021), holds promise for enablement of high throughput measurements of *tln* and *cobleaf.* Furthermore, deep learning and image processing approaches to extract tassel flowering patterns (Mirnezami et al., 2021)could potentially enable the parameterization of models aimed at predicting hybrid maize production.

The MFS discussed in this manuscript is appropriate for non-stress field conditions, as seed production fields are managed to avoid water and nitrogen deficit, but an interesting extension of our work would be to integrate water stress, to be able to predict and simulate genotype specific traits in both stress and favorable field conditions. Indeed, in case of water stress, extrusion can be delayed, silk receptivity can be reduced (Khan et al., 2022), and the number of pollen grains can be altered (Hall et al., 1982). In addition, water stress slows down ear biomass accumulation (Edmeades et al., 1993). This would allow us to broaden the scope of this type of work by being able to increase the range of environmental conditions to sample additional levels of genotypes and could help exploring future climate change scenarios to optimize breeding strategies.

## Conclusion

The MFS-WGP infrastructure, integrating whole genome prediction with the MFS crop growth model can accurately predict unobserved physiological parameters and simulate maize flowering phenotypes, as part of in-field in-season inbred growth development. It does so by leveraging genetic, environmental and management data in an integrated mathematical model that is scalable and easy to use. An example of an important practical application of this method is the ability to recommend differential planting intervals for male and female maize inbreds used in commercial seed production fields to synchronize male and female flowering.

## Data Availability statement

The data that support the findings of this study are not publicly available.

## Conflict of Interest statement

All authors declare that they have no conflicts of interest.

## Author Contributions

AL: conceptualization, data curation, formal analysis, visualization, methodology, investigation, writing-original draft, editing

EM: conceptualization, formal analysis, methodology, investigation, writing-review and editing

HZ: conceptualization, software, formal analysis, methodology, visualization, investigation, writing-review and editing

FT: conceptualization, formal analysis, methodology, investigation, writing-review and editing

EW: data collection, writing-review and editing RC: conceptualization, methodology

JP: data collection, writing-review and editing

RT: conceptualization, formal analysis, methodology, investigation, writing-review and editing, supervision

## Abbreviations

ASI: anthesis interval silking
CGM: crop growth model
coblf: coefficient of leaf appearance
ebR1: ear biomass at reproductive stage R1
egr: ear growth rate
GDD: growing degree day
LN: leaf number
MAE: mean absolute error
MFS: maize flowering synchrony
RMSE: root mean square error
tln: total leaf number
WGP: whole genome prediction

## Supplemental materials

### S1. Additional information about weather characteristics of the estimation set

Meteorological conditions differed between growing seasons (Figure S1). Mean air temperature during early vegetative growth was higher for year 2018 (20.8 oC) as compared to year 2020 and 2021 (14.8 and 16.1 oC, respectively). Mean air temperature during late vegetative stage was higher for year 2021 (24.1 oC) as compared to year 2018 and year 2020 (22.9 and 21.7 oC, respectively). Accumulated rainfall during late vegetative stage was approximately 50% lower in year 2021 than in the other two years.

**Figure S1.**
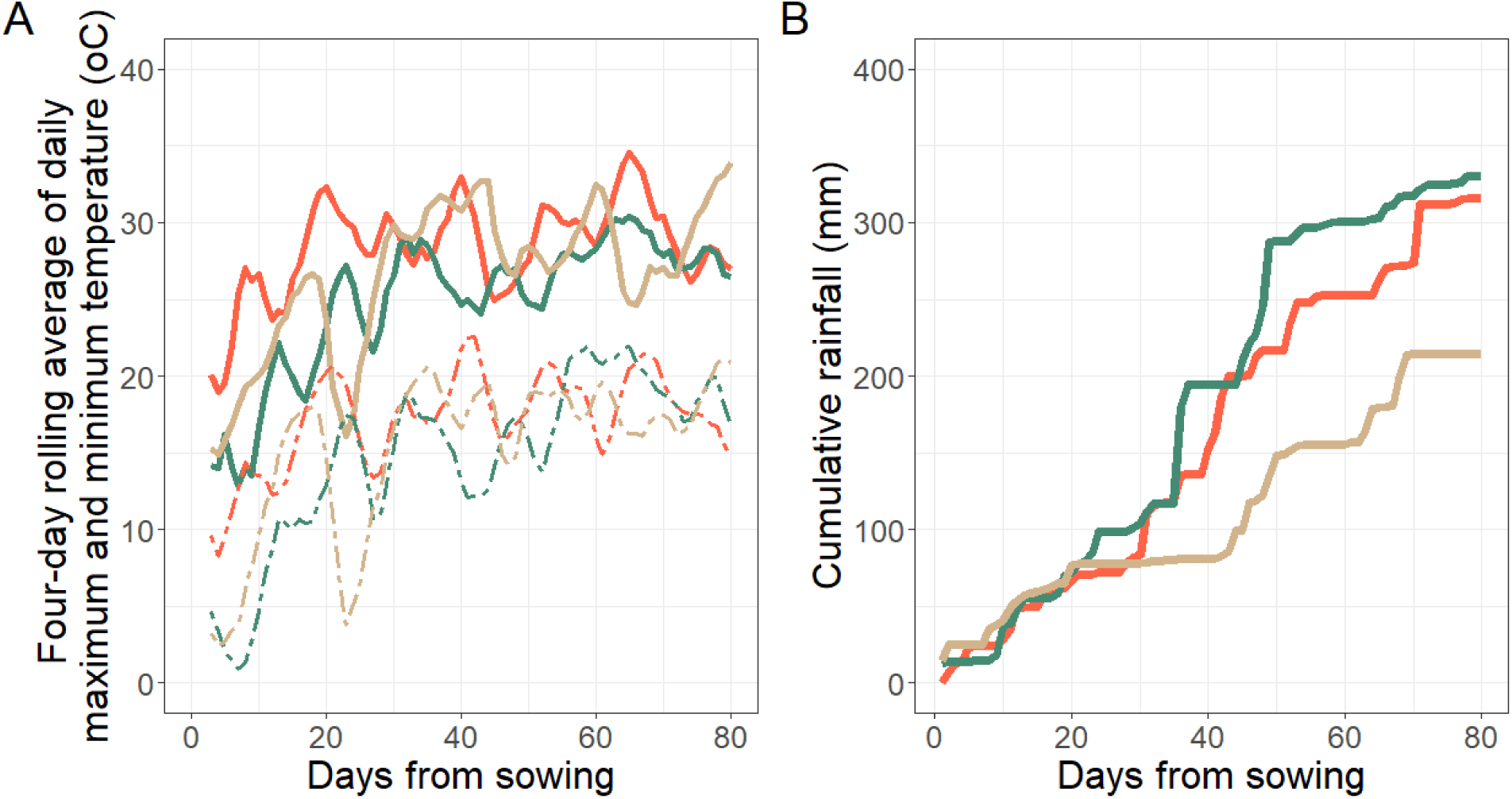
(A) Maximum and minimum air temperature evolution (solid and dashed line, respectively) in three growing seasons (Year 2018: red; Year 2020: green and Year 2021: beige), and (B) cumulative rainfall in millimeters. Data are presented as a function of days from sowing.

### S2. Impact of Uninformative priors on predicted physiological parameters and simulated phenotypes

To stay consistent with the two set of priors described in the manuscript, only the prior for ebR1 was set as a weak prior with a mean and standard deviation chosen such that the 2.5 and 97.5 quantiles range from approximatively 0 to 50, which reflects that negative values are biologically impossible but otherwise doesn’t restrict the value range of the posterior.

The priors for ebR1 (called here *ebR1_weak*), *tln* and *coblf* are described as follows:

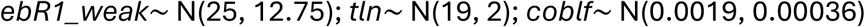

**Figure S1a:**
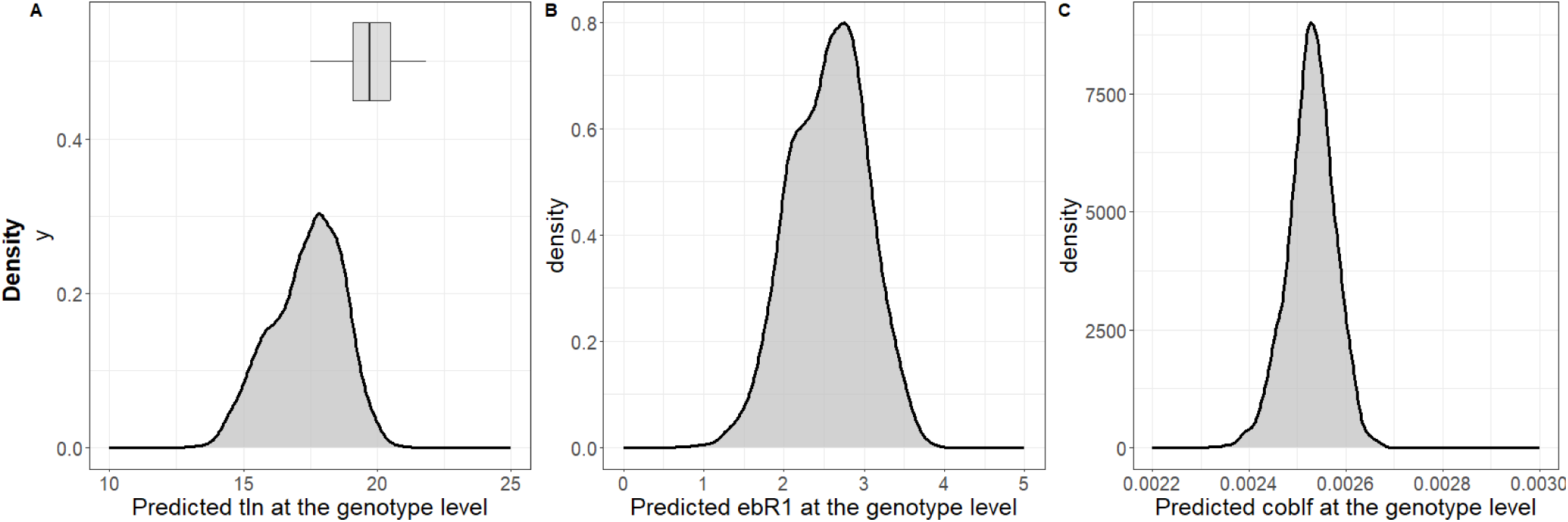
Distribution of the posterior mean of *T_ti_* for *tln* (A), *ebR1*_*weak* (B), and *coblf* (C) of the ∼2000 female lines in the data set A

**Figure S2:**
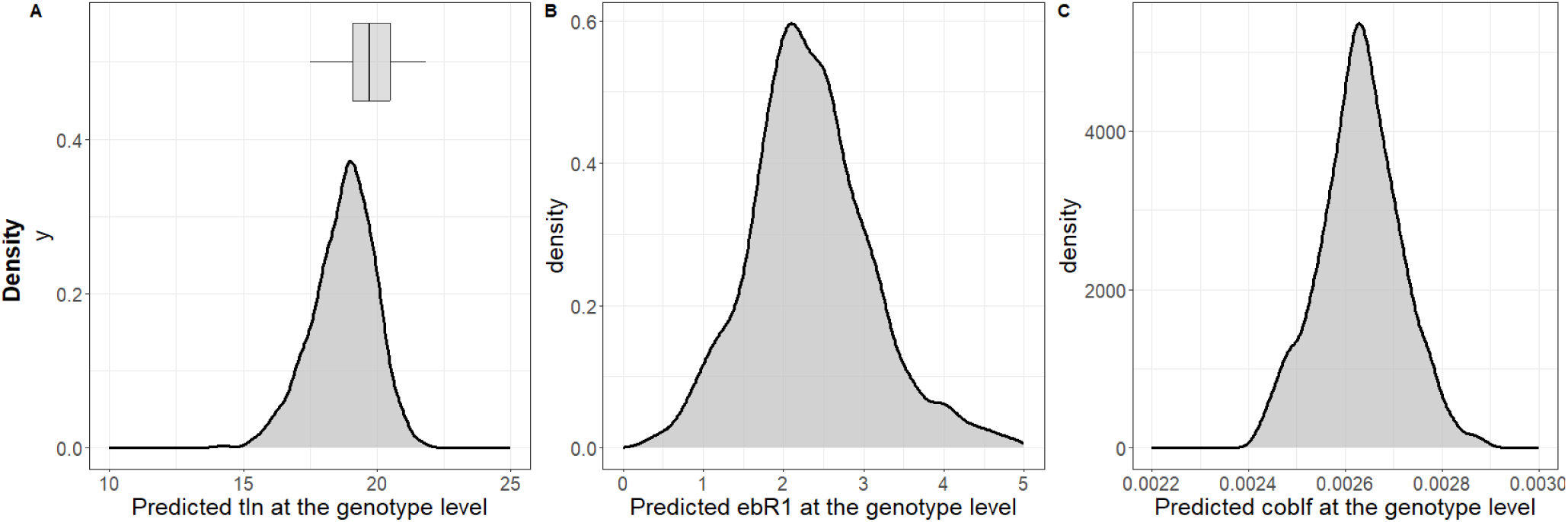
Distribution of the posterior mean of *T_ti_* for *tln* (A), *ebR1_weak* (B), and *coblf* (C) of the ∼500 female lines in the data set B.

The distribution of the posterior mean of *T_ti_* for *tln* does not fall into the distribution of observed *tln* (grey boxplot Figure S1A and S2A) whatever the size of the estimation set. When the estimation set is small, the distribution of *ebR1_weak* covers values (0 to 5 grams) that are not realistic compared to observed ear biomass at R1 stage as observed in the literature (see section 2.1 in the manuscript).

**Figure S3:**
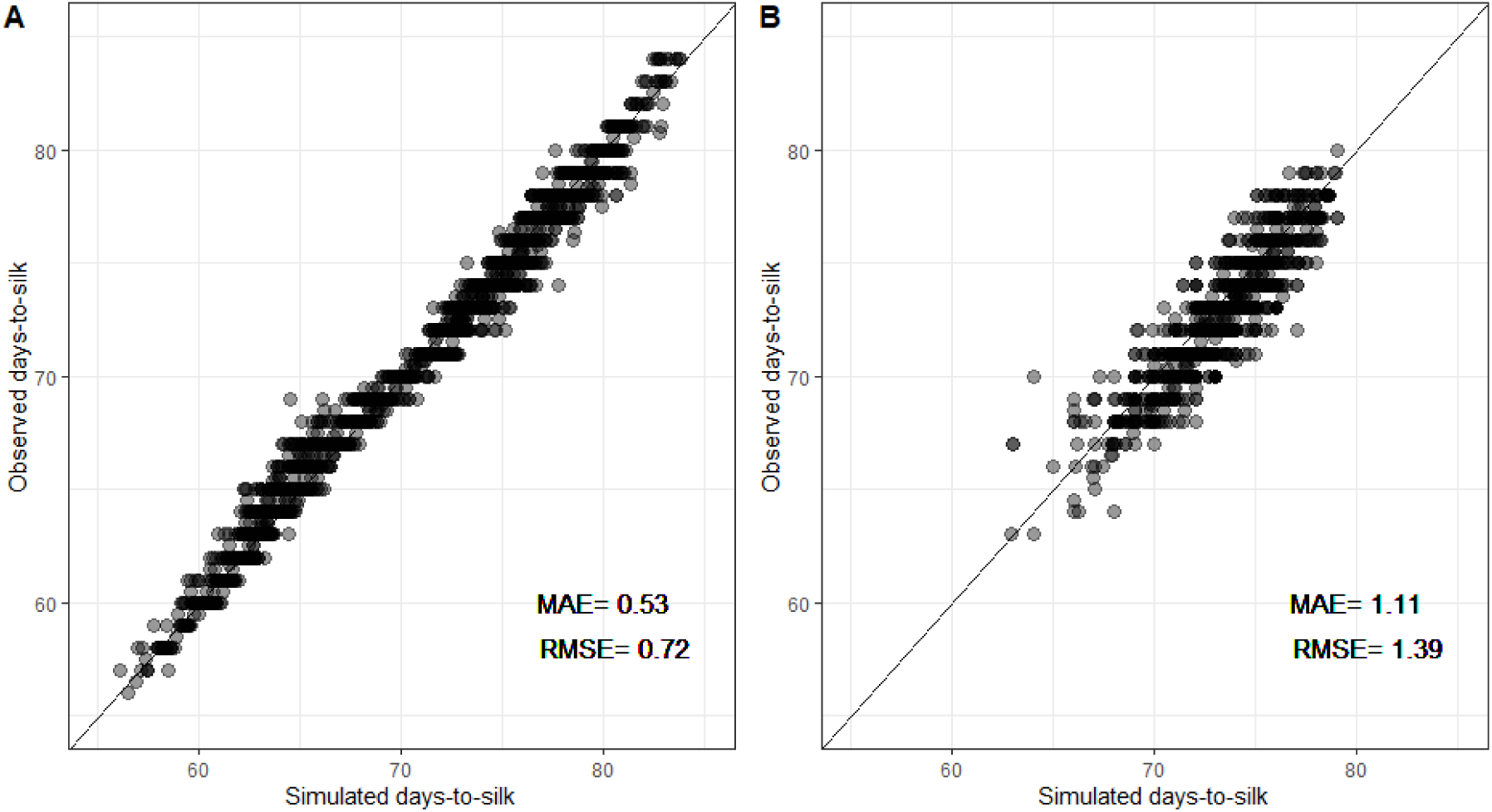
Simulated days-to-silk compared to the observed value by data set A (large data set of ∼2000 genotypes, on the left (A)) and data set B (smaller data set of ∼500 genotypes, on the left (B)) for a weak prior *ebR1_weak*. The mean absolute error (MAE) and root mean square error (RMSE) are indicated at the bottom right of each plot. The identity line is indicated in black.

When the data set is large and diverse in terms of fields and genotypes (Figure S3A), there is no impact of the weak prior on the simulated phenotypes (MAE = 0.53 and RMSE = 0.72). On the opposite, when the data set is small and restricted to one field and only ∼16% of the genotypes (Figure S3B), there is a strong impact of the prior chosen as shown by the MAE and the RMSE (MAE equals 1.11 and RMSE equals 1.39).

As a reminder, assessing the impact of weak priors on the predicted physiological parameters and simulated phenotypes was presented here as a baseline. Using weak or uninformative priors open the risk to create improper distributions as showed by the posterior mean of *T_ti_*(Figure S1-S2) while informative priors relfects existing knowledge about physiological parameters.

## Notes

### Competing Interest Statement

The authors have declared no competing interest.

